# Serum metabolic signatures of cognitive resilience in a longitudinal aging cohort

**DOI:** 10.64898/2026.03.29.715122

**Authors:** Toon A.W. Scheurink, Jeong In Seo, Lurian Caetano David, Crystal X. Wang, Deyvis Solis, Jasmine Zemlin, Jaclyn Bergstrom, Pieter C. Dorrestein, Ipsita Mohanty, Anthony J.A. Molina

## Abstract

Aging is typically accompanied by a progressive decline in cognitive function, yet some individuals maintain exceptional cognitive performance, even across the transition from middle to older age, defining exceptional cognitive resilience. While existing measures of resilience primarily rely on clinical assessments, its molecular determinants and early predictive markers remain poorly understood. Here, we performed untargeted LC-MS/MS profiling of longitudinal serum samples to identify metabolic signatures associated with cognitive resilience, which was established based on cognitive tests conducted over 28 years in a cohort of 237 participants. We observed associations across multiple chemical classes, including carnitines, glutamine conjugates, phosphocholines, as well as diet-and drug-derived metabolites. Chemical class-specific analyses revealed distinct metabolic profiles, including predominantly negative associations of medium-chain acylcarnitines with cognitive resilience, increased accumulation of glucuronide conjugates in individuals with low cognitive resilience, altered metabolism of the antihypertensive drug, metoprolol, and elevated levels of dietary compounds such as piperine and lutein in individuals with high cognitive resilience. By leveraging public metabolomics data, we further contextualized the metabolic signatures with respect to their organ specificity, microbial origin, and disease associations. Collectively, these metabolic features, including several previously underexplored compounds, represent promising candidates for functional characterization in mechanisms of aging biology and provide mechanistic insights into the molecular basis of cognitive resilience.

**Highlights:**

- Serum metabolite MS/MS features, including acyl carnitines, phosphocholines, and hippuric acid conjugates, are associated with cognitive resilience in 237 individuals transitioning from middle to older age.
- Diet-derived piperine is positively associated with cognitive resilience.
- Differences in β-blocker drug metabolism, rather than parent drug levels, are associated with cognitive resilience.
- Repository-scale searches for the resilience-associated metabolites reveal organ specificity, microbial contributions, and their presence across multiple disease contexts.

**Graphical abstract:** 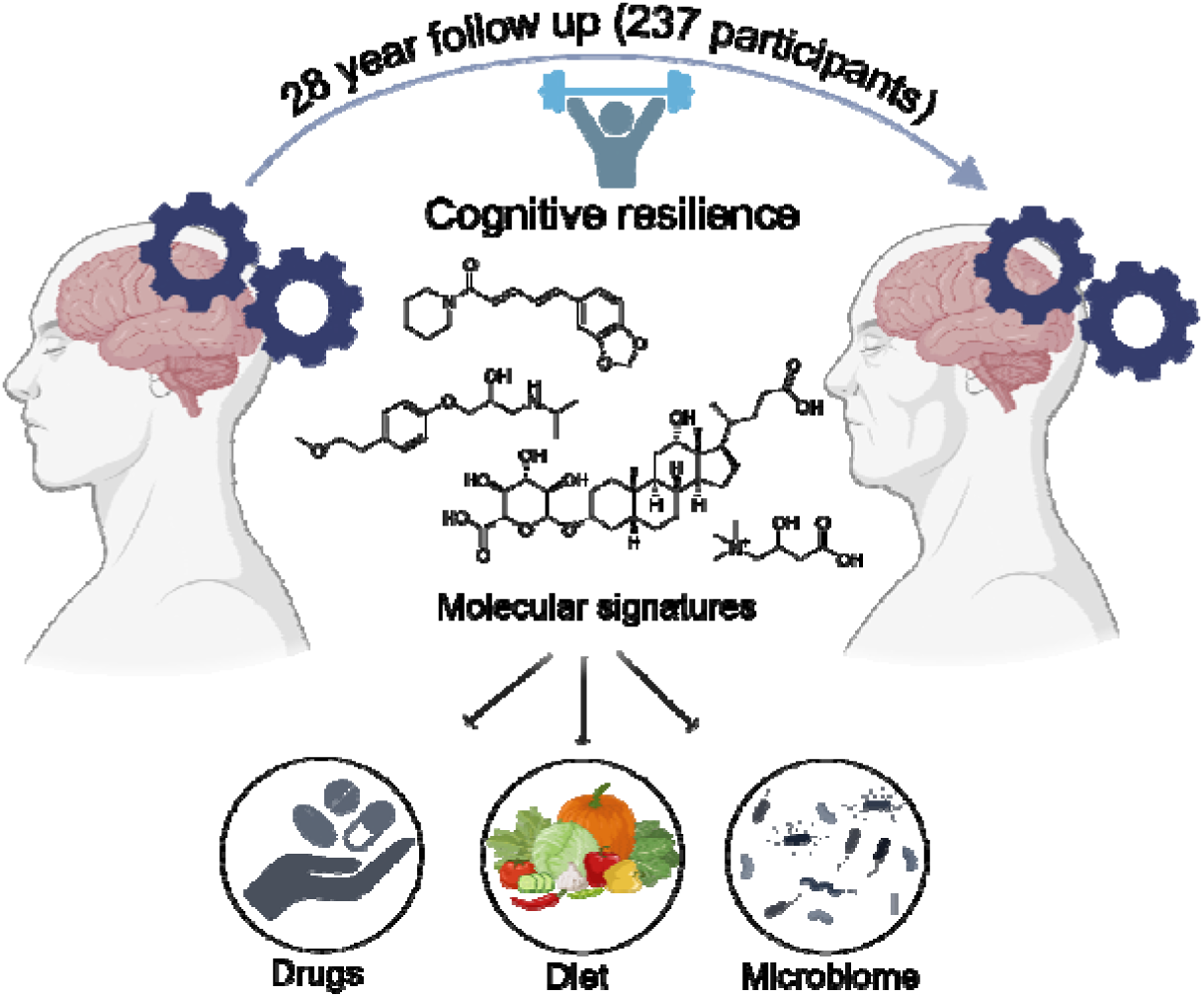

## Introduction

Typical age-related changes in cognitive performance are well documented across numerous longitudinal cohort studies, and are apparent even among individuals who do not develop neurodegenerative disease.^1^ Importantly, heterogeneity in the pace and severity of cognitive decline is also widely recognized. However, the molecular mechanisms that shape the trajectory of aging remain poorly understood. As life expectancy increases, the number of individuals living with age-related cognitive impairment and dementia continues to rise, placing a substantial burden on caregivers, health systems, and societies worldwide.^2^

Interestingly, previous studies have demonstrated that some older adults are able to maintain normal cognition even in the presence of significant neuropathological burden, including amyloid plaques, tau tangles, or mixed pathologies. In contrast, others exhibit cognitive impairment even with comparatively mild pathology.^3–6^ These differences highlight the definition of cognitive resilience, which refers to the ability to maintain cognitive performance despite physiological and pathological changes that accumulate with aging.^7,8^ Although lifestyle factors such as education, physical activity, and vascular health contribute to resilience, these influences explain only part of the inter-individual variability, and the biological bases of resilience are currently not well understood.^2,7^ Emerging evidence points to several biological domains that may underlie resilience. These include preserved synaptic and neuronal plasticity, efficient mitochondrial and bioenergetic function, and regulation of neuroimmune and inflammatory responses.^9^ Such processes likely interact across multiple systems, motivating interest in systems-level molecular approaches for characterizing resilience. Mass spectrometry-based metabolomics is a powerful tool for investigating biochemical pathways associated with aging, particularly in cognitive impairment.^10,11^ Systematic reviews and targeted metabolomic analyses consistently show alterations in lipid metabolism, including phosphatidylcholines, sphingomyelins, and other membrane-associated lipids, as well as changes in amino acids, redox metabolites, and energy substrates among individuals with Mild Cognitive Impairment (MCI) or Alzheimer’s Disease (AD).^9,12–16^ Plasma metabolomic trajectories linked to concurrent cognitive and mobility decline suggest that lipid and amino-acid pathways are sensitive indicators of future dementia.^17^ Despite these advances in disease states, no large-scale metabolomics study has yet defined the molecular signatures of aging or cognitive resilience phenotypes. As a result, the molecular foundations that enable certain individuals to resist cognitive decline remain largely unknown.

Our study aims to address this gap by applying untargeted Liquid Chromatography-Tandem Mass spectrometry (LC-MS/MS)-based serum metabolomics in a deeply phenotyped cohort of 237 older adults to identify metabolic signatures associated with aging in general and cognitive resilience. Metabolomics analysis aids in understanding these biochemical pathways, provides insight into mechanisms supporting preserved cognition in aging, and informs strategies for promoting healthy cognitive longevity. Moreover, these lay the foundation for predictive biomarkers of cognitive health, which can support the advancement of proactive healthcare strategies.

## Results

### Cohort description and analysis workflow

Serum samples for metabolomics analysis were obtained from 237 participants in the Rancho Bernardo Study (RBS) of Healthy Aging. RBS is a population-based, longitudinal study with over 6,726 participants followed for up to 47 years. ^18,19^ It has extensive metadata, including clinical, cognitive and demographic outcomes (**Figure 1a**). For the purpose of this study, we utilized the cognitive resilience scores, calculated based on a participant’s performance on several cognitive tests, representing different modalities of cognitive ability, over time in a subset of 2,616 participants (*see methods for calculation of resilience scores*). The cognitive scores were calculated from tests conducted over a period of 28 years. Each participant is assigned a single score for the entire study, reflecting their trajectory over time. A lower cognitive resilience score indicates greater decline in cognitive function, whereas a higher score indicates a more stable cognitive function across study visits. Detailed information on the calculation of the cognitive resilience scores can be found in a previous publication.^20^

**Figure 1.**
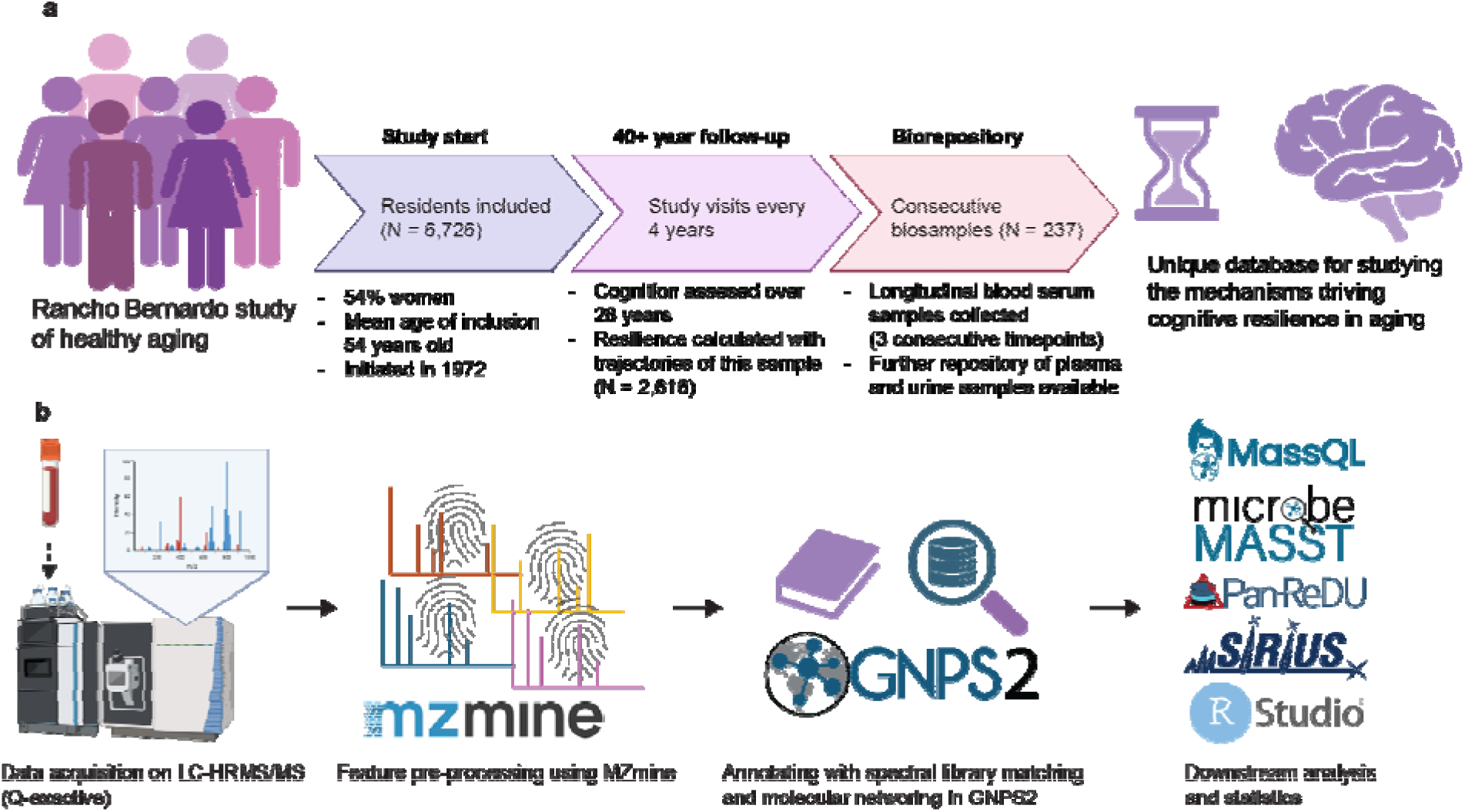
Study setup and workflow. **a)** The Rancho Bernardo study setup. **b)** Sample processing and feature detection methods employed, as well as the utilized downstream analysis tools. Icons were obtained from Biorender, logos from their respective websites.

For metabolomics analysis, we utilized fasting serum samples collected at 3 consecutive time points within RBS for 237 participants. The serum samples were analyzed using untargeted LC-MS/MS-based metabolomics. The complete workflow for the metabolomics analysis is shown in **Figure 1b**. We aimed to investigate the molecular underpinnings of aging in general and then for cognitive resilience without confounding by age. Therefore, for cognitive resilience, we selected 1 visit per participant that resulted in the lowest standard deviation in age, thereby harmonizing age effects across individuals. The characteristics of the selected cohort for the cognitive resilience analysis is highlighted in **Table 1**.

**Table 1.** Characteristics of the participants from the RBS cohort included in the metabolomics analysis for cognitive resilience.

| Participant characteristics | N = 237<sup>1</sup> |
| --- | --- |
| Age | 73.8 (3.0) |
| Sex |  |
| female | 152 (64%) |
| male | 85 (36%) |
| Cognitive resilience score | 0.2 (0.4) |
| Selected Visit |  |
| V5 | 41 (17%) |
| V8 | 55 (23%) |
| V10 | 83 (35%) |
| V12 | 58 (24%) |

<sup>1</sup>Mean (SD); n (%)

### Linear Mixed Modeling Detects Stable MS/MS Features Associated with Aging

First, we aimed to find metabolites that were consistently associated with chronological age within participants. We performed this analysis on all 711 samples (all 3 visits from each participant). After sparsity selection, we obtained 562 features that were present in <70% of samples. Here, the term feature refers to a chromatographic peak characterized by a mass-to-charge ratio (*m/z*) and retention time pair and quantified by integrated peak area following MZmine preprocessing.^21^ We then performed a linear mixed model and selected features with a heterogeneous slope across individuals. 113 features were significantly and singularly associated with age, which are presented in **Supplementary Table 1**. Notably, most of these 113 features (90.3%) were positively associated with age, indicating a broad age-related shift in the metabolomics profile.

### sPLS Regression Identifies Stable MS/MS Features Associated with Cognitive Resilience

Next, to identify metabolite features associated with cognitive resilience, we performed sparse partial least squares regression (sPLS-R) on the untargeted LC-MS/MS data. Optimal sparsity parameters were determined using a ranked scree-style plot and cross-validation, resulting in the selection of 3080 features and 1 component, indicating that a single multivariate dimension best captured the association between metabolomic features and cognitive resilience. The model explained 20% of the total variance in cognitive resilience (p < 0.001) (**Figure 2a**). Observed R^2^ using 10-fold cross-validation (cv) was 0.148 (root mean square error (RMSE) = 0.41). The selected features showed high selection stability, with a median stability score of 1.0 (range: 0.2-1.0), with 92% of features selected in at least 7 of the 10 cross-validation folds (**Supplementary Figure S1**). From a total of 22,753 features, our model selected 3080 (13.5%) features (**Figure 2b**). The 3080 features, their weight, stability and slope over time can be found in **Supplementary Table 2**. We aimed to provide metabolite class information for the 3080 features associated with cognitive resilience using SIRIUS.^22^ Of these 3080 model-selected features, 2098 could be assigned to chemical classes using SIRIUS through the CANOPUS (Class Assignment and Ontology Prediction Using mass Spectrometry) module. Prediction and classification of the features were performed using NPClassifier for biologically relevant molecules, with superclasses (NPC) corresponding to broad metabolite classes associated with major biosynthetic pathways.^23,24^ The pie chart (**Figure 2c**) depicts the annotation of 1,053 features to their matching NPC superclass with over 70% probability. Four major superclasses (i.e., glycerophospholipids, small peptides, fatty esters, and oligopeptides) represent over 50% of the total counts for the matching 56 superclasses.

**Figure 2.**
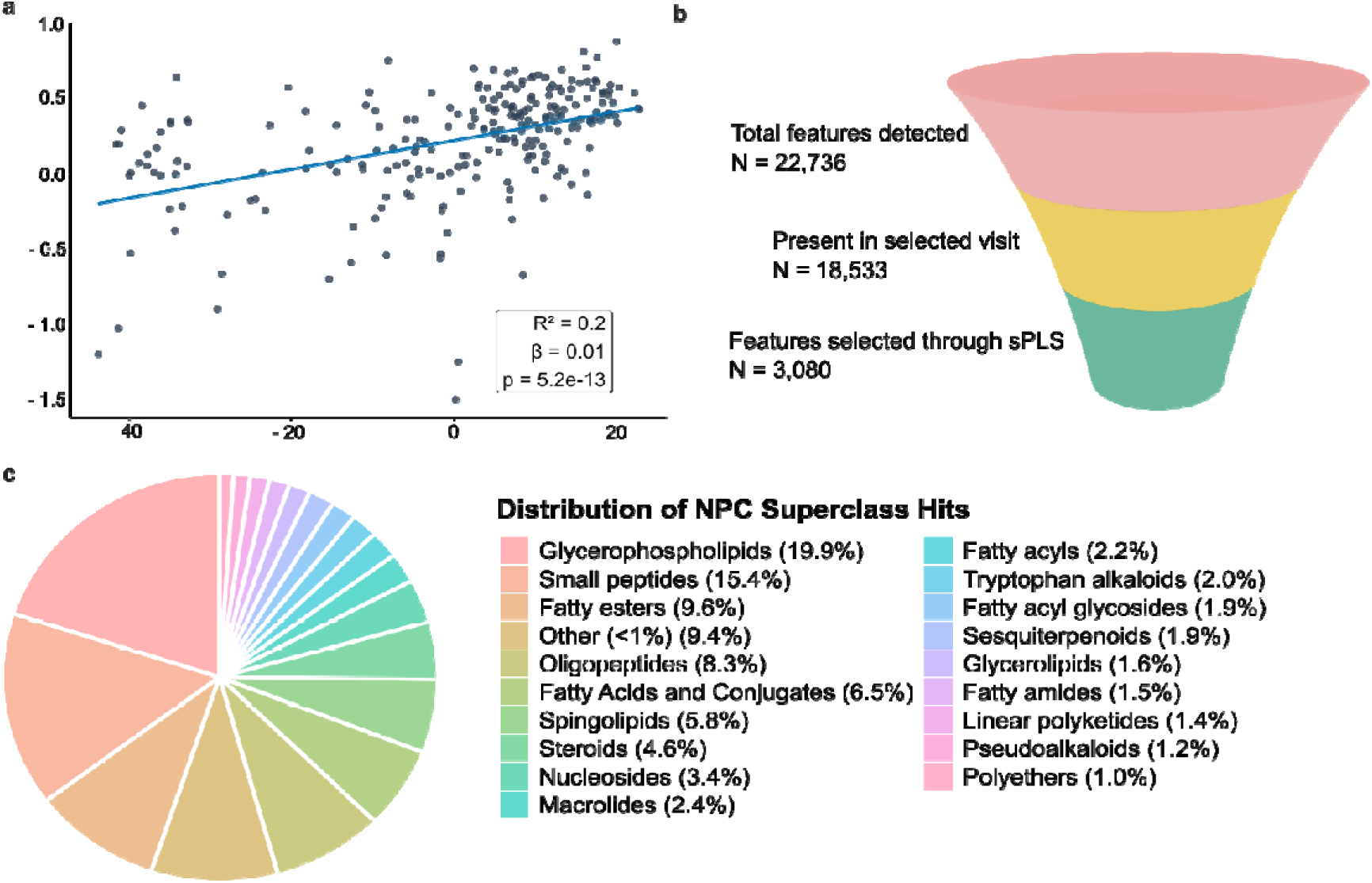
Model performance and class of cognitive resilience-associated features. **a)** A linear regression of the first component of the sPLS model against the cognitive resilience scores. **b)** Selection of the 3080 features associated with cognitive resilience. **c)** Distribution of different chemical classes within the selected 3080 features, determined by SIRIUS 6.3.2.^22^ Icons were obtained from Biorender.

Next, we validated our model for feature selection using an external visit per participant that was not used in the initial model training, chosen to be closest in age to the selected visit. This longitudinal validation allowed us to assess which features remained robust and informative over the time course. The external validation model could significantly predict variation in cognitive resilience based on the selected features (R^2^ = 0.02, p = 0.024, 10-fold CV observed r = 0.165). When sPLS-R was applied to the external sample selection using identical parameters, the selected metabolite feature sets showed moderate overlap (Jaccard similarity = 0.471) and consistent effect directions (loading weight correlation = 0.542). Notably, features with higher weights in the original model were substantially more stable, with 92% of metabolites in the top 5% by weight reselected in the external analysis, compared with an overall feature overlap of 64% (**Supplementary Figure S2**).

To investigate longitudinal age-related changes in the 3080 selected features, we used linear mixed-effects models (LMM) to assess their associations with age across all three study visits. To accommodate the LMM, we selected features that were present in at least 40% of the samples, resulting in 974 features. Out of these, 733 were (BH-corrected) and non-singularly associated with age. Interestingly, there was a positive association between weight in the sPLS model and the slope of features over time (R^2^ = 0.57, L = 3.8, p < 2e-16) (**Supplementary Figure S3**). Together, these results indicate that features with higher predictive weight for cognitive resilience also showed more consistent longitudinal age-associated behavior. Also, the quadrant plot shows that only 19 of 113 aging-related metabolites were not associated with cognitive resilience, while the remaining metabolites showed varying patterns of association with both aging and cognitive resilience (**Supplementary Figure S4a**). These patterns included positive correlations with both aging and cognitive resilience, as well as positive associations with aging but negative associations with cognitive resilience. Furthermore, chemical class annotation of the aging-associated metabolite features was performed using SIRIUS, which identified three major compound classes among the selected aging-related metabolites (i.e., fatty acyls, oligopeptides, and small peptides) (**Supplementary Figure S4b**).

### Molecular Networking Highlights Chemical Families Associated with Cognitive Resilience

To visualize the molecular families and their association with positive or negative cognitive resilience trajectory scores, we used molecular networking (**Figure 3**).^25^ The molecular network visualizes different chemical families detected in this dataset based on MS/MS spectral similarity. Each circular node in the molecular network represents a molecular feature, the node size reflects relative abundance based on peak area, and the node color indicates the direction of association with cognitive resilience. Molecular features that share structure similarity and consequently spectral similarity are connected by an edge in the same subnetworks. On inspection of the molecular network, we observed that the largest and most interconnected cluster (highlighted in green in **Figure 3**) contains nodes with MS/MS matches to acylcarnitines, spanning short (C2-C5), medium (C6-C12), long (C13-C21), and very long (≥C22) chain fatty acids.^26^ Further, we obtained spectral matches to the newly discovered *N*-acylcarnitines and some candidate carnitines, where the exact chain length and the modification are not yet characterized.^27^ Overall, 83% of the carnitines captured in this dataset were negatively associated with cognitive resilience. The prominence of these carnitines suggests substantial remodeling of fatty-acid oxidation pathways across samples, consistent with known roles of acylcarnitines in energy metabolism, mitochondrial function, and metabolic stress.^26^

**Figure 3:**
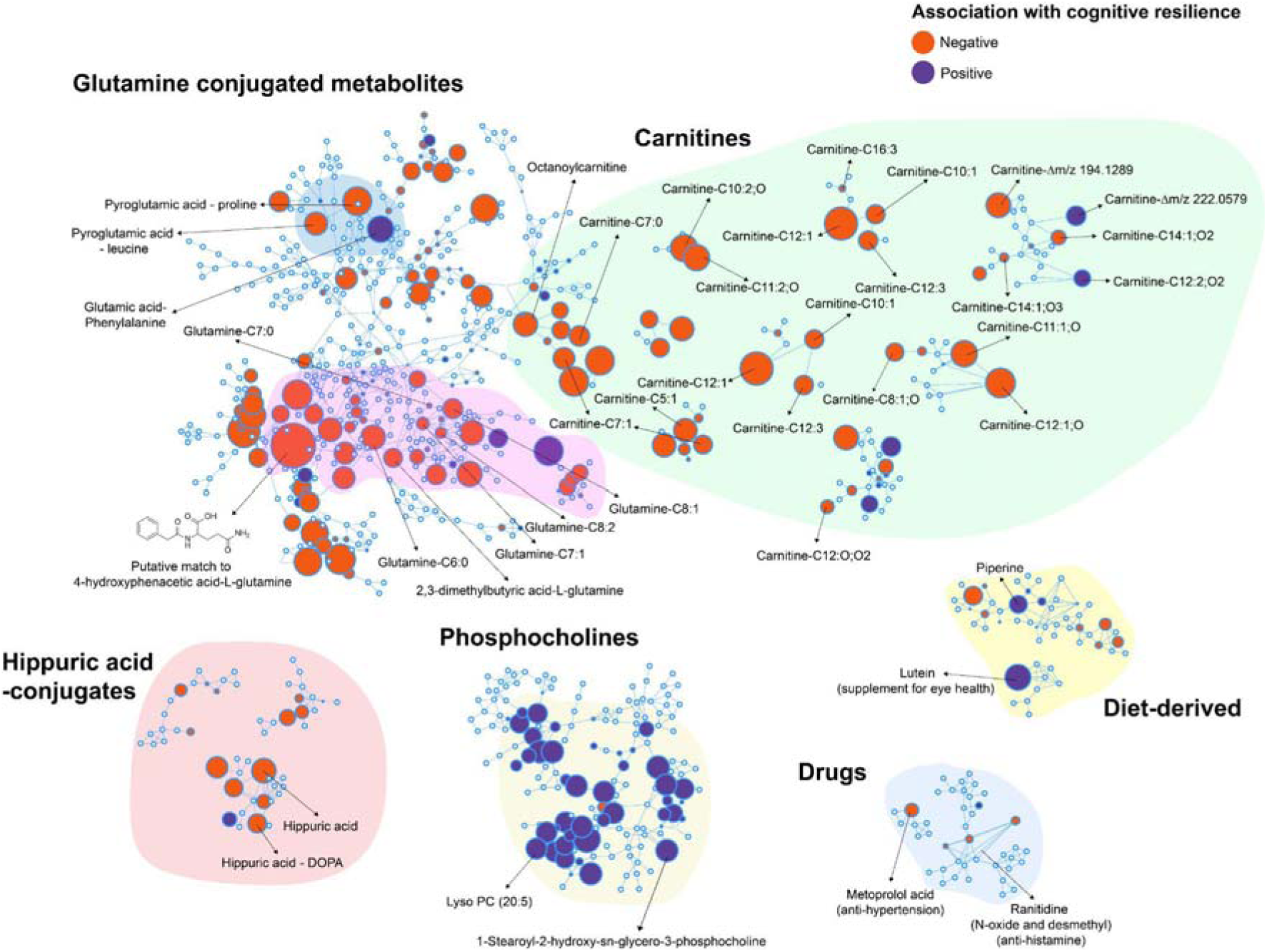
M**o**lecular **Network of the features associated with cognitive-resilience:** A subset of molecular families is shown and highlighted from the complete molecular network. Nodes represent individual molecular features and are colored blue or red according to the direction of association with cognitive resilience. Node size reflects the weight of the feature on the component in the sPLS-R model.

In addition to the carnitines, another major subnetwork of metabolites negatively associated with cognitive resilience was that of the glutamine-conjugated metabolites. The annotated metabolite features within this cluster were those of glutamine conjugated fatty acids (C6:0, C7:0, C7:1, C8:1, and C8:2). In addition, two other metabolites within this cluster were those of a putative match to 4-hydroxyphenacetic acid conjugated with glutamine and 2,3-dimethylbutyric acid conjugated to glutamine. Positively loaded on the component associated with cognitive resilience was a cluster of phosphocholines, molecules essential for cell membranes. These were sparsely annotated, but included 2 lysophosphatidylcholines. Two other metabolites with strong negative associations with cognitive resilience were pyroglutamic acid conjugated to proline and leucine, which were also observed in the same cluster as the glutamine conjugated metabolites. These observations were interesting in the context of recent flux analysis studies showing up to 30% increase in glutamine flux in aging mice.^28^ Finally, a smaller cluster of hippuric acid-related conjugates was associated negatively with cognitive resilience.

### Acylcarnitines, Bile Acids, and Glucuronides Are Associated with Cognitive Resilience

As discussed in the previous section, many compounds within specific chemical groups were identified to be associated with cognitive resilience based on the sPLS-R model. In particular, carnitines with diverse structural variations (i.e., acylcarnitines), which formed the largest molecular family in our metabolomics analysis, were especially notable. To further investigate their associations with cognitive resilience, we performed a hurdle model analysis for carnitines included among the 3,080 features selected from the sPLS-R model (**Supplementary Table 3**). In the hurdle model, we first detect whether the absence or presence of a feature is associated with cognitive resilience. We then leave out samples where we do not detect that feature, and test if the abundance of a feature is related to cognitive resilience. From this analysis, we identified 69 carnitines with statistically significant correlations based on presence versus absence (FDR < 0.05) (**Figure 4a**). Only seven of these showed positive correlations with cognitive resilience. In the samples where they were detected, 83 carnitines were significantly associated with cognitive resilience (FDR < 0.05) (**Figure 4c**). All of these were negatively associated with cognitive resilience. In both steps, most of these carnitines were medium chain length acylcarnitines, ranging from C6 to C12, based on putative annotations by spectral library matching in GNPS2 (**Figure 4b and 4d**; **Supplementary Table 3**).^29^ In addition, specific acylcarnitines that demonstrated negative association (C12:1 carnitine, both in presence versus absence and in abundance models; **Figure 4e**) or positive association (C9:0 carnitine, only in presence versus absence model; **Figure 4f**) showed significantly different peak area patterns in individuals with high or low cognitive resilience.

**Figure 4.**
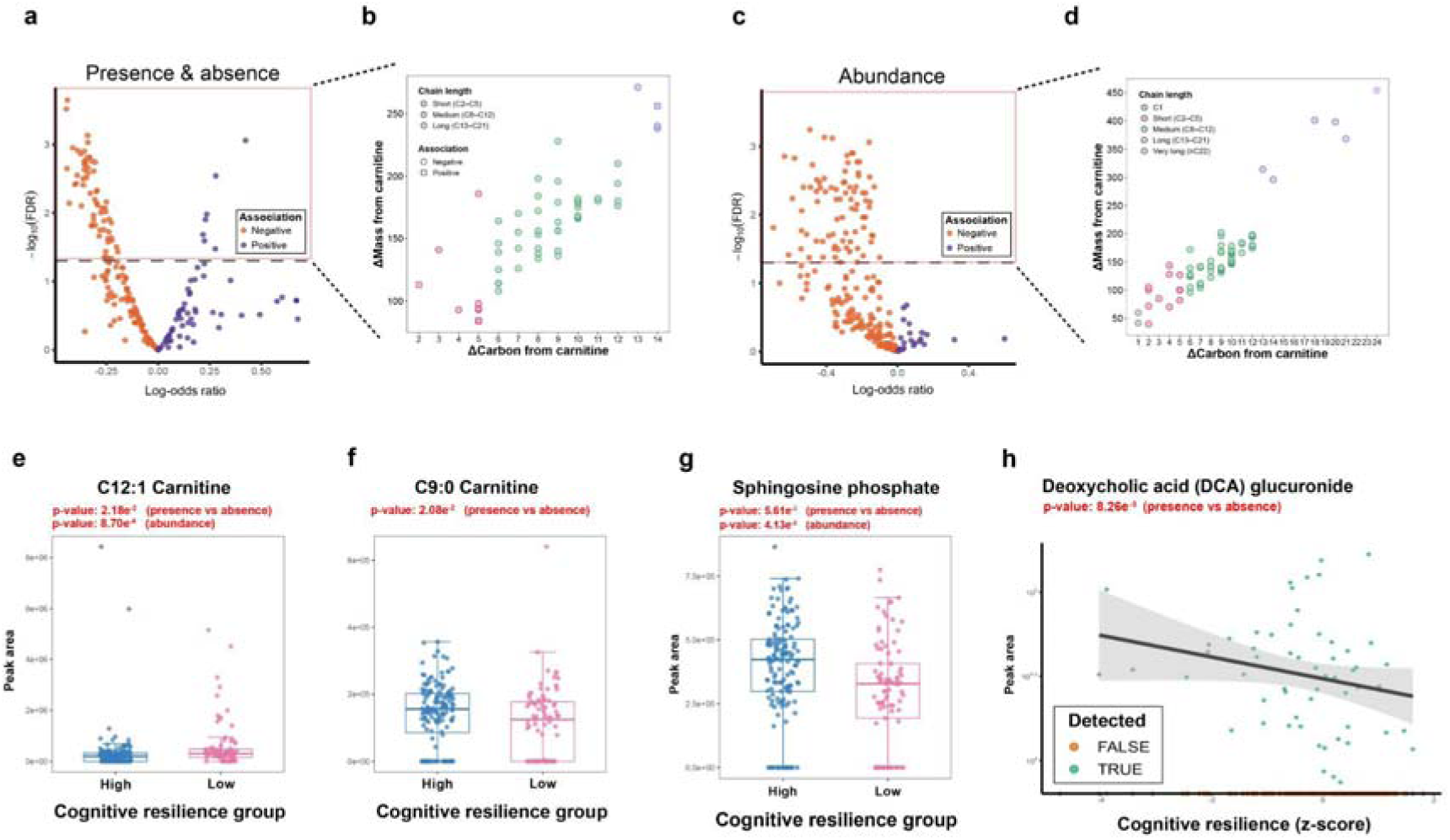
Carnitines, sphingolipids and glucuronides are associated with cognitive resilience. **a)** First step of the hurdle model, a presence vs absence plot showing the positive or negative correlations of acylcarnitines with cognitive resilience. **b)** Chain length distribution of carnitines showing statistical significance in the first step of the hurdle model (FDR < 0.05). **c)** Second step of the hurdle model, showing if the abundance of carnitines in samples where they are detected is associated with cognitive resilience. **d)** Chain length distribution of carnitines showing statistical significance in the second step of the hurdle model (FDR < 0.05). **e)** Peak area abundances of C12:1 carnitine and **f)** C9:0 carnitine in individuals with high or low cognitive resilience. **g)** Peak area distributions of sphingosine phosphate in individuals with high or low cognitive resilience. **h)** Linear regression plot of deoxycholic acid glucuronide. Statistical differences were evaluated using a hurdle model, and p-values were adjusted for multiple comparisons using the Benjamini-Hochberg method. Compounds were putatively annotated based on spectral library matching in GNPS2. Individuals were classified into high or low cognitive resilience groups based on quartiles of cognitive resilience scores within the complete RBS cohort, with the lower two quartiles assigned to the low group and the upper two quartiles assigned to the high group.

From the overall molecular network, we identified sphingolipids as generally positively associated with cognitive resilience. We plotted the peak area of a specific lipid mediator, sphingosine phosphate, which showed significantly higher levels in individuals with high cognitive resilience in both the presence/absence and abundance models (**Figure 4g**). In addition, the bile acid deoxycholic acid (DCA) glucuronide showed a significant negative association with cognitive resilience in the presence/absence model (**Figure 4h**). Glucuronidation is a major detoxification pathway, catalyzed by UDP-glucuronosyltransferase (UGT) enzymes, which are widely expressed in the liver and extrahepatic tissues.^30^ Given this broad distribution, we next sought to systematically identify additional glucuronide conjugates associated with cognitive resilience using the Mass Spectrometry Query Language (MassQL), a domain-specific language that enables concise and flexible querying of mass spectrometry data.^31^ We designed the MassQL query to capture the characteristic neutral losses of glucuronide conjugates, specifically 176.0321 Da and 194.0425 Da, in the positive ion mode.^32^ This MassQL-based analysis enabled us to identify 129 potential glucuronide conjugates among the 3,080 features previously selected by the sPLS-R model, and 81.4% of these showed negative correlations with cognitive resilience in the sPLS component (**Supplementary Figure S5a**). Fold change analysis further revealed significant group differences in their peak areas, with most glucuronide conjugates observed at higher levels in individuals with low cognitive resilience (**Supplementary Figure S5b**).

### Diet-and Drug-Related Metabolites Associated with Cognitive Resilience

In addition to the carnitines and glucuronidated metabolites described above, we also identified metabolites originating from dietary sources or pharmaceutical compounds that were significantly associated with cognitive resilience. Notably, two diet-derived compounds were identified. Piperine, a major bioactive compound derived from black pepper, was significantly more often present in individuals with high cognitive resilience (**Supplementary Figure S6a**). Similarly, lutein, a phytochemical known for its role in supporting visual function and commonly obtained from green leafy vegetables or dietary supplements,^33,34^ was more prevalent in individuals with high cognitive resilience and was significantly more often present in those with high cognitive resilience (**Supplementary Figure S6b**).

Interestingly, we identified a metabolite of the antihypertensive drug metoprolol, metoprolol acid, in the molecular network, which had a negative correlation with cognitive resilience (**Figure 3**). Meanwhile, the parent drug metoprolol and its structural analogue atenolol, both belonging to the same class of antihypertensive drugs, β-blockers, were also detected in our dataset through spectral library matching. Neither parent drug, metoprolol or atenolol, showed a significant correlation with cognitive resilience, suggesting that variations in how individuals metabolize these compounds may be associated with cognitive resilience. Hence, we sought to identify additional metabolites of metoprolol and atenolol that may be associated with cognitive resilience using MassQL query analysis, as drug metabolites often preserve much of the parent drug structure and therefore generate similar MS/MS spectra.^35^ Given the structural similarity between metoprolol and atenolol (**Figure 5a**), which share a common aryloxypropanolamine backbone (highlighted in **Figure 5a**) but differ in their side chains, with an acetamide group in atenolol and a methoxy group in metoprolol, we designed a MassQL query targeting shared fragment ions derived from the aryloxypropanolamine structure (*m/z* 72.0806, 98.0964, 116.1070, and 133.0643). The MassQL filtering yielded 46 MS/MS spectra for putative metoprolol or atenolol metabolite ion variants. To reduce potential false discoveries, only MS/MS detected together with their respective parent drugs in the same samples were retained after co-occurrence filtering. Metabolites in the co-occurrence plot (**Figure 5b**) were putatively annotated based on rationalization of chemical formulae corresponding to the mass shifts from parent drug and known drug metabolic pathways, together with a MassQL query targeting metabolites from glucuronidation (**Supplementary Figure S7a**). The unannotated compounds were represented only by their delta mass relative to each parent drug. Using this approach, we observed metoprolol modifications consistent with glucuronidation, hydroxylation, and demethylation, as well as combinations of these processes, such as hydroxylation with glucuronidation or demethylation (**Supplementary Figure S7b)**. In addition, an atenolol metabolite generated through glucuronidation was detected (**Figure 5b**). Furthermore, these compounds were verified as metabolites of the parent drugs based on their earlier retention times relative to the parent drugs (**Supplementary Figure S7c and S7d**). This analysis confirmed that the metabolites of atenolol/metaprolol were not different ion forms/adducts of the parent drugs but were real metabolized products. Notably, comparison of metoprolol and its metabolite levels between the low and high cognitive resilience groups revealed distinct metabolic patterns (**Figure 5c**). Individuals with high cognitive resilience exhibited higher levels of metabolites from glucuronidation, whereas those with low cognitive resilience showed higher levels of metabolites from oxidation, such as hydroxylated metabolites.

**Figure 5.**
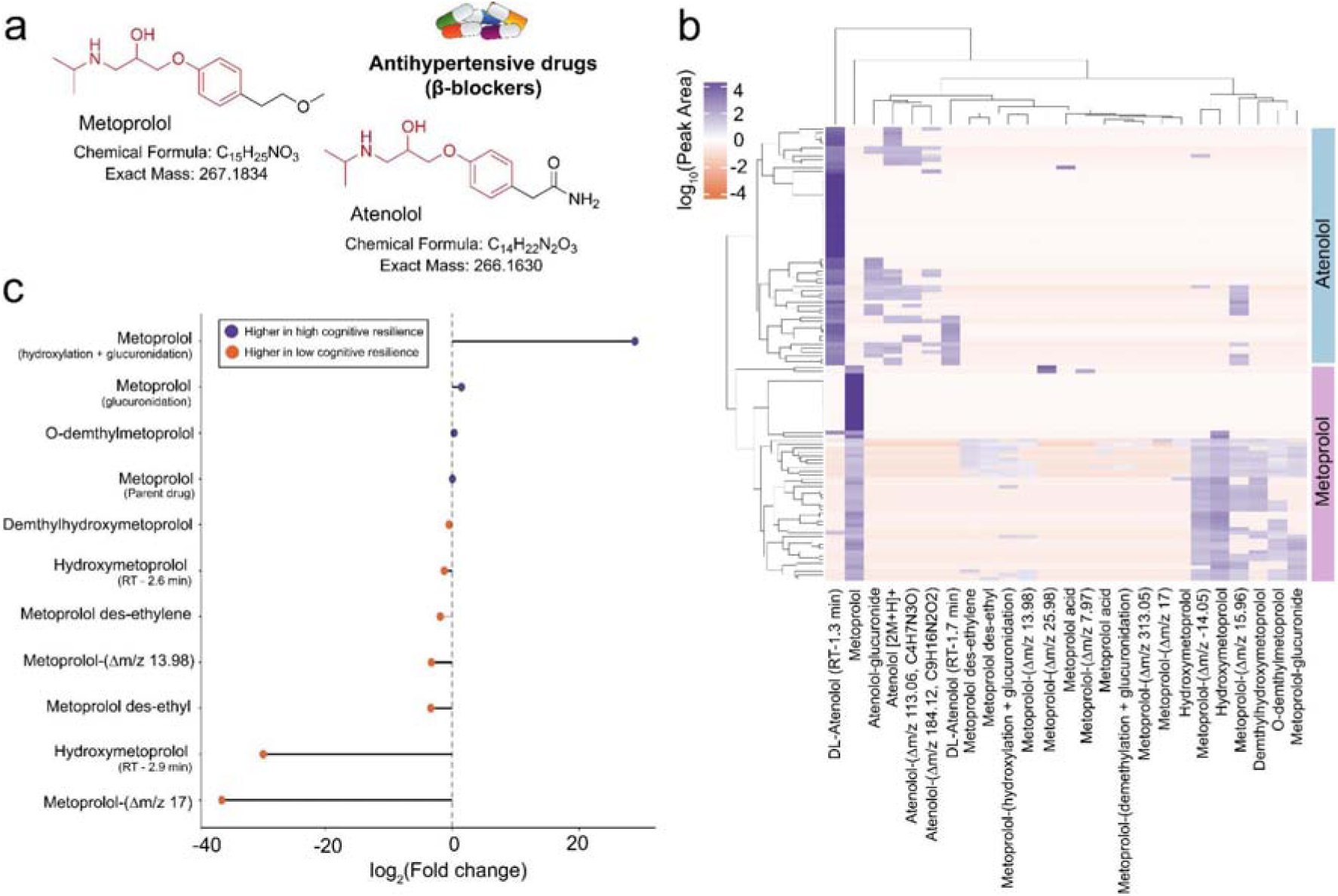
Patterns of association of antihypertensive drug metabolites with cognitive resilience. **a)** Chemical structures of metoprolol and atenolol (the common aryloxypropanolamine backbone is highlighted in red). **b)** Co-occurrence plot mapping putative metabolites of metoprolol and atenolol. **c)** Fold change analysis comparing metoprolol metabolite profiles between the low and high cognitive resilience groups. The icon was obtained from <u>Bioicons.com</u>.

### Repository-Scale Spectral Searches Reveal Tissue, Disease, and Microbial Contexts of Cognitive Resilience-Associated Features

We sought to provide broad contextualization for the 3080 cognitive resilience-associated metabolites based on their distribution across public metabolomics repositories. Therefore, we used MS/MS spectral matching with the Mass spectrometry search tool (MASST)^27^ to identify the distribution patterns of these features across different tissue matrices, organisms, and health conditions. MASST enables searching tandem mass spectra against curated reference databases, including GNPS/MassIVE, MetaboLights, and Metabolomics Workbench.^29,36,37^ In addition, integration with the Pan-Reanalysis of Data User Interface (Pan-ReDU)^28,29^ provides access to harmonized and curated metadata linked to these repositories, enabling systematic contextualization of matched spectra.

Of the 3,080 features with MS/MS, 2,626 (85.3%) were matched to other datasets in public repositories. In total, these matched 4,094,433 MS/MS spectra from 354,952 unique files in 3,119 datasets. We used the Uber-anatomy ontology (UBERON)^38^ classification in Pan-ReDU to explore the distribution of features with positive or negative loadings on the cognitive resilience-associated component (**Figure 6a**). Features matched differentially to tissue matrices, with matches to lung, heart and epithelial cell samples likely to be positively loaded, whereas features matched to urine, kidney and skin were more likely to be negatively loaded. Interestingly, features found in the brain were slightly more likely to be positively associated with cognitive resilience, whereas features matching cerebrospinal fluid were often negatively associated. Most of our samples available in the pan-ReDU datasets were blood plasma and serum, followed by fecal samples (**Figure 6b**).

**Figure 6.**
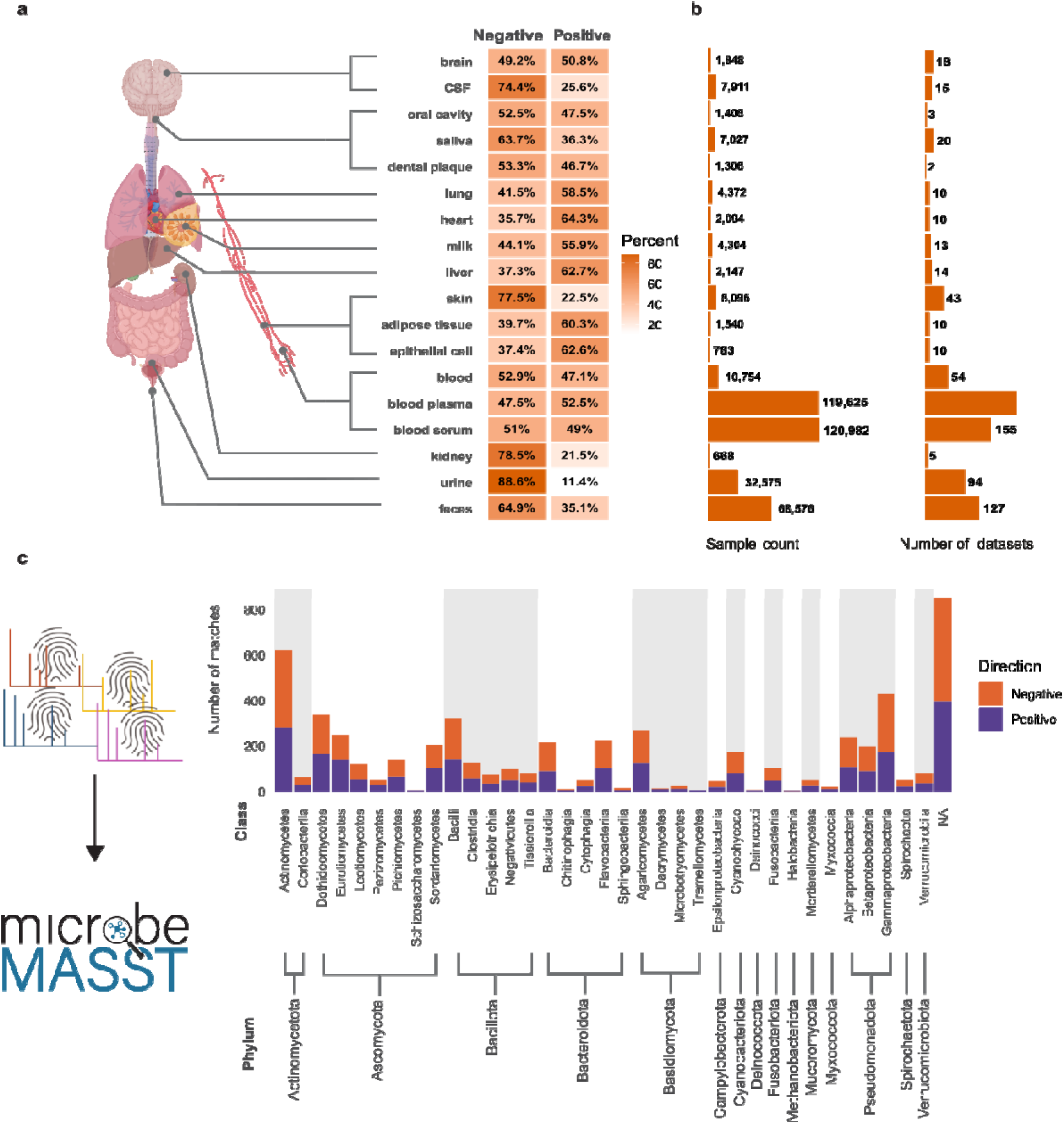
**Repository-scale contextualization of cognitive resilience-associated features across tissues and microbial cultures**. **a)** Distribution of features matched to human studies across different tissue matrices using MASST and Pan-ReDU metadata. **b)** The availability of these samples in the ReDU datasets. **c)** Count and taxonomic distribution of features matched exclusively to microbial monocultures in external libraries, summarized by class and phylum. CSF - cerebrospinal fluid. Icons were obtained from Biorender, and logos were obtained from the respective websites.

Next, we determined whether the 3,080 selected spectra matched spectra from human samples with different health conditions. To prevent confounding effects of sample type on the distribution of positive and negative features, we included only blood samples (plasma and serum) when matching spectra to health conditions. These samples that are most prevalent in the pan-ReDU datasets are osteoarthritis, Alzheimer’s disease and Chagas disease. Matches to AD and osteoarthritis were more frequently negative. The features that matched samples from HIV and (history of) AIDS cohorts were mostly positively loaded on the component associated with cognitive resilience, although this was based on 112 and 118 samples, respectively (**Supplementary figure S8**).

In addition, we used the domain-specific MASST, microbeMASST, to identify features that were detected in microbial monocultures.^39^ Out of the 3080 features, 1100 (35.7%) matched to microbeMASST. We obtained a total of 230,645 matches to MS/MS spectra from 19,474 unique files across 313 datasets. MicrobeMASST also contains human cell line cultures. By excluding blanks, QC’s and scans that match to human cell lines, we determined that 742 scans were found in microbial monocultures. Using National Center for Biotechnology Information (NCBI) taxonomic classifiers, we found that these features matched microbial monocultures spanning six kingdoms, 15 phyla, 35 classes, 94 orders, 217 families, 399 genera, and 967 species across the domains Archaea, Bacteria, and Eukaryota. The highest number of matches were obtained for microbial phyla such as actinomycetes and gammaproteobacteria. Although there were no clear patterns in whether those features were positively or negatively associated with cognitive resilience (**Figure 6c**).

To evaluate the association between microbial-derived features and cognitive resilience, we performed PLS regression using only the 720 features identified from microbial monocultures. This microbial-restricted model explained 17% of the total variance in cognitive resilience (p < 0.001), only slightly lower than the full model. Ten-fold cross-validation yielded an observed R² of 0.146 (RMSE = 0.341). We then replicated the same external validation strategy used for the full model. Notably, the microbial-restricted model showed improved external predictive performance, significantly explaining variation in cognitive resilience (R² = 0.04, p = 0.0011; 10-fold cross-validated r = 0.21). This analysis demonstrates that features detected in microbial monocultures retain predictive power for cognitive resilience even when considered independently of the full metabolomics feature set.

## Discussion

This study provides untargeted LC-MS/MS serum metabolomics profiles across 237 participants, linked to aging and cognitive performance trajectories measured over up to 28 years in the Rancho Bernardo Study of Healthy Aging. For aging in general, we identified 113 metabolite features that showed consistent changes with chronological age across participants, as well as a significant association at the population level. Moreover, we identified 3,080 metabolite features that individually contribute to cognitive resilience and provide examples of how levels of selected molecules, including carnitines, phosphocholines and drugs change with cognitive resilience and also their association with aging in general. By leveraging MS/MS spectral matching to public datasets in metabolomics repositories,^40^ we show how these metabolite features are distributed across different tissue types, diseases and also if they are detected in microbial monocultures.

We show that metabolomics signatures are associated with cognitive resilience and validate that model performance and many of the (especially high-weight) identified features remain stable across different sampling times within participants. Prevalent positive resilience-associated features increase as participants age, whereas negative resilience-associated features decrease or disappear with increasing age. Using feature-based molecular networking, we observed that metabolite features clustered into broad chemical classes within shared subnetworks. Notably, features within the same subnetworks tended to associate with cognitive resilience in the same direction, suggesting coordinated behavior of metabolites arising from related biochemical pathways. A network of carnitines, molecules involved in energy production, is consistently associated negatively with cognitive resilience. This may reflect decreased mitochondrial function^41^, and age-related increase in levels of acylcarnitines have previously been identified.^42^ Indeed, mitochondrial function as measured in peripheral mononuclear blood cells was lower in people with dementia compared to healthy controls, and associated positively with cognitive ability.^43^ In contrast, phosphocholines, molecules essential for cell membranes, were near-consistently associated positively with cognitive resilience.^44^ They are precursors to and degradation products of phosphatidylcholine, which promotes functional neurogenesis.^44,45^ The sphingosine phosphate, a phosphocholine, is a key signaling lipid that regulates diverse pathophysiological processes across multiple organ systems, including the central nervous, cardiovascular, and immune systems.^37^ Notably, reduced sphingosine phosphate levels in the human brain have been linked to the pathogenesis of Alzheimer’s disease,^38^ and its receptor is a therapeutic target in neurology, as demonstrated by the approval of Fingolimod for relapsing multiple sclerosis.^39^ In our metabolomics analysis, sphingosine phosphate levels were significantly higher in individuals with high cognitive resilience (**Figure 4g**), suggesting that loss of sphingosine phosphate may be more common in those with low cognitive resilience.

In addition, a network of glutamine-conjugated molecules was associated mostly negatively with cognitive resilience. Glutamine is one of the most common precursors for glutamate, the most abundant excitatory neurotransmitter.^46^ Underactive glutamate transmission has been associated with cognitive impairment, and dietary supplementation of glutamine has been shown to delay cognitive decline in mice.^47^ However, both acute and chronic excesses of glutamate may lead to neurotoxicity and neurodegeneration.^48^ Within this context, the glutamine-conjugated molecules captured in our cognitive resilience model may reflect glutamine-glutamate dysregulation in cognitive impairment. Additionally, we identified a metabolite derived from glucuronidation with significant negative correlation with cognitive resilience (**Figure 4h**), which is a well-recognized detoxification process mediated by UDP-glucuronosyltransferase (UGT) enzymes.^30^ This process prevents the accumulation of potentially harmful molecules in the body. Thus, the observed correlation may reflect greater accumulation of the parent compounds in individuals with low cognitive resilience. This glucuronidated metabolite, DCA glucuronide, is a microbial bile acid. Bile acids are important signaling molecules regulating metabolic, immune, and neurophysiological processes, and their chemically diverse forms, shaped by host and microbial metabolism, remain underappreciated.^49–53^ Accordingly, elevated DCA glucuronide may reflect alterations in gut microbial composition and bile acid signaling linked to cognitive function. Supporting this interpretation, previous studies have reported higher concentrations of secondary bile acids, including DCA, in the plasma of individuals with cognitive impairment.^54^ These metabolomic patterns related to detoxification pathways suggest that cognitive resilience may be associated with a reduced burden of potentially toxic compounds, which would otherwise require conjugation and elimination through metabolic processes such as glucuronidation.

In addition, several diet-or drug-derived metabolite signatures associated with cognitive resilience were observed. One of the diet-related metabolites we observed with positive association with cognitive resilience was piperine. Although evidence for piperine’s therapeutic relevance in human cognitive impairment remains limited, several animal studies have reported its effectiveness in preventing cognitive impairment in Alzheimer’s disease models.^55^ These effects have been suggested to involve mechanisms such as inhibition of necroptosis, as reported in a separate study.^56^ From a pharmacokinetic perspective, a large proportion of intact piperine can circulate in the bloodstream due to its relatively low susceptibility to hepatic metabolism, and piperine levels in the brain have been reported to be comparable to those in plasma.^57^ Taken together, higher levels of piperine in individuals with high cognitive resilience may contribute to the prevention of cognitive decline. Furthermore, given previous findings linking visual decline to cognitive decline during aging^58^ or in neurological disorders such as dementia,^59^ our observation of higher levels of lutein may contribute to the prevention of age related cognitive decline. Lutein is a carotenoid that comes almost entirely from plant-based foods, especially leafy greens, and is protective for retinal function and linked to eye health.^33^

We next observed distinct metabolic patterns of the antihypertensive drug metoprolol associated with cognitive resilience. The observed patterns suggest that individuals with higher cognitive resilience may preferentially convert metoprolol to more readily cleared glucuronide conjugates, whereas individuals with lower cognitive resilience accumulate metabolites arising from oxidative metabolism (**Figure 5b**). Although studies examining the relationship between cognitive decline and drug metabolism are scarce, one study demonstrated that glucuronidation of paracetamol was selectively reduced in frail elderly individuals, while glucuronidation did not differ between healthy young adults and healthy elderly subjects.^60^ This suggests that glucuronidation may decline only under specific health conditions, such as frailty. By contrast, glucuronidation appears to be largely preserved with aging, as consistently reported in other studies.^61,62^ Hence, the higher levels of metoprolol glucuronides observed in individuals with high cognitive resilience may indicate a potential link between cognitive resilience and glucuronidation activity.

The metabolite features associated with cognitive resilience are distributed differentially across public LC-MS/MS datasets. First, we identified that features associated positively or negatively are differently distributed among different tissue matrices. For example, the majority of matches to kidney and urine was to features negatively associated with cognitive resilience. This may reflect the filtration function the kidneys perform, with harmful features removed from the circulation. Indeed, cognitive decline is common in people with chronic kidney disease, which has been partly attributed to accumulation of uraemic toxins and metabolic dysregulation.^63^ Additionally, these results emphasize the importance of tissue matrix selection for future studies on cognitive resilience, and advocate for a variety of sample types in a study when possible. Second, we found that features associated positively or negatively are additionally differently distributed among different disease types. Notably, the features matching to samples from people with Alzheimer’s disease were predominantly associated negatively with cognitive resilience. In contrast, features matching to samples from people with HIV or with history of AIDS were mostly positively associated with cognitive resilience. In general, risk factors of HIV-associated cognitive impairment are multisystemic, and exacerbated by substance use and mental health comorbidities as well as social or environmental risk factors (e.g., poverty, homelessness).^64,65^ Therefore, while the neuroprotective features may be similar across cohorts, neurotoxic features may differ due to discrepant risk factors between people with HIV and our cohort of predominantly white older adults from high socioeconomic backgrounds.

Using microbeMASST,^39^ we identified that 742 out of the 3,080 features were found in microbial but not human monocultures. These are likely related to the gut microbiome, which is known to exert effects on cognition and more through the gut-brain axis.^66^ On their own, these features associated well with cognitive resilience, indicating that microbial metabolites are impactful for cognitive function in later life. Despite that, across microbial classes and phyla, we did not observe any group that was consistently associated with features either positively or negatively related to cognitive resilience. This may be due to some features matching to multiple microbe monocultures, diluting the effects. Future studies combining metabolomics with metagenomics may provide more depth for exploring how specific microbes and their metabolites can affect long-term brain health.

In conclusion, this study provides the first detailed metabolomics-based overview of metabolite features associated with cognitive resilience, emphasizing the importance of particular chemical classes including previously uncharacterized carnitines and phosphocholines. Untargeted metabolomics studies on datasets focused specifically on cognitive resilience are scarce. Here, we address this gap by systematically parsing metabolomic signatures linked to long-term cognitive outcomes. Our results demonstrate that metabolomic profiles are associated with cognitive resilience, suggesting that these features may relate to ongoing or future cognitive decline. This dataset provides a resource for follow-up experimental and longitudinal studies to determine which metabolites may contribute to preserving or impairing cognitive function over time.

### Limitations of the study

This study should be considered in light of its limitations. Firstly, metabolite annotations are performed based on MS/MS spectral matches with GNPS2 libraries (level 2-3).^29^ Further validation of the MS/MS matches to get accurate structures with stereochemistry and regiochemistry defined will require retention time and/or ion mobility matching with synthetic standards. A substantial fraction of detected features could not be annotated through spectral library matching. Many molecules have no MS/MS characterization, and differences in instrumentation and fragmentation means that even known molecules may not match to spectral libraries.^67^ Similarly, spectral matching with MASST tools is dictated by the available datasets in public repositories. As more datasets are added, future reanalysis of this data may provide new and extended insights. Second, the metabolomics signature provided here is based on the Rancho Bernardo Study (RBS) of Healthy aging, which is a relatively homogenous cohort in terms of ethnicity and socioeconomic status, limiting generalizability while reducing confounders.^19^ Despite these limitations, we have shown that untargeted LC-MS/MS metabolomics can be used to provide an expansive view of the metabolites associated with cognitive resilience and aid the understanding of the complex, dynamic metabolic processes underlying cognitive resilience during aging.

## Resource availability

### Lead contact

Further information and requests for resources and reagents should be directed to the lead contact, Anthony Molina.

### Materials availability

This study did not generate unique reagents.

### Data and code availability

The untargeted LC-MS/MS data is publicly available on GNPS/Massive under the accession MSV000097935. The corresponding GNPS2 job is available at https://gnps2.org/status?task=6ea14530712c4b1ba27f638961b76c24. Code and scripts used for analysis in this manuscript is available on GitHub: https://github.com/ToonScheurink/Resilient_aging_metabolomics/tree/main/Scripts.

## Acknowledgements

This work was funded by The Wellcome Leap Dynamic Resilience program (co-funded by Temasek Trust). Data collection for the Rancho Bernardo Study of Healthy Aging was provided primarily by the National Institutes of Health (including grant numbers: HV012160, AA021187, AG028507, AG007181, DK31801, HL034591, HS06726 and HL089622).

Archiving and sharing of RBS data was supported by the National Institute on Aging: AG054067. J.I.S. is supported by the National Research Foundation of Korea (NRF) (RS-2025-02373133). L.C.D. is supported by the Fulbright U.S. Student Program through the DDRA (Doctoral Dissertation Research Award), which is sponsored by the U.S. Department of State and the Fulbright Brazil Commission.

## Author contributions

Conceptualization, I.M., P.C.D., A.M.; methodology, P.C.D., I.M., T.S., J.I.S.; formal analysis, I.M., T.S., J.I.S., L.C.D.; investigation, I.M., T.S., J.I.S., L.C.D., C.W., D.S., J.Z., J.B.; resources, J.B., A.M.; writing - original draft, I.M., P.C.D., A.M., T.S., J.I.S., D.S.; writing - review & editing, all authors; supervision, I.M., P.C.D., funding acquisition, P.C.D., A.M.

## Declaration of interests

P.C.D. is an advisor and holds equity in Cybele, Sirenas, and BileOmix, and he is a scientific co-founder, advisor, and holds equity and/or receives income from Ometa, Enveda, and Arome with prior approval by UC San Diego. P.C.D. consulted for DSM Animal Health in 2023.

## Declaration of generative AI and AI-assisted technologies in the writing process

During the preparation of this work, the authors used ChatGPT/Gemini to improve text flow, clarity, simplify technical language, and perform grammar checks. After using this tool/service, the authors reviewed and edited the content as needed and take full responsibility for the content of the publication.

## Methods

### Study population

Samples were collected from a subset of 237 participants in the Rancho Bernardo Study of Healthy Aging (RBS).^19^ This is a prospective cohort study initiated in 1972 including residents of 30+ years old in Rancho Bernardo, San Diego, California. At inclusion, the cohort totaled 6,726 participants, representing 82% of the adults. These community dwellers were followed up approximately every 4 years starting in 1984, collecting a wide range of detailed demographic, health and biosample data. There were no exclusion criteria. All study visits were approved by the University of California San Diego Human Research Protections Program (UCSD IRB 960019; UCSD IRB 040165; UCSD IRB 130757). All participants provided written informed consent at each clinic visit.

### Calculation of cognitive resilience score

Cognitive function was assessed in 2,672 participants starting at visit 5 (performed between 1988-1992) and subsequent visits. Participants performed a battery of tests, including the Trail-Making Test part B (psychomotor tracking and executive function), the Mini Mental State Exam (global cognitive function), the Category Fluency Test: animal naming (semantic memory and executive functioning) and the Buschke-Fuld Selective Reminding test (verbal episodic memory). A linear mixed-effects regression model based on these combined test z-scores could be built for 2,616 participants, including fixed effects (age, sex and education) and random participant-level intercepts and slopes. Each participant’s resilience score is the difference between their person-specific slope and the overall model slope. The overall model slope was-0.7, indicative of a-0.7 SD decrease in combined test score per decade after 70 years of age.

### Sample collection and preparation

Blood samples were obtained via venipuncture between 07:30 and 11:00 after a requested 12 hour fast. Serum was separated and frozen at-70 C upon collection in sunlight-protected tubes. We included 237 participants of 50+ years old who had cognitive resilience scores and had sufficient available samples from three consecutive study visits covering a period of at least 15 years. Metabolomics analysis was performed on these 711 serum samples. Serum samples were processed using the Phenomenex Phree™ 96-well phospholipid removal kit (Phenomenex Inc., Torrance, CA, USA). For each sample, 100 µL of serum was carefully pipetted onto the center of the filter membrane in each well. To maintain a 4:1 (v/v) solvent-to-sample ratio, 400 µL of 100% methanol was added directly to each well containing serum to facilitate protein precipitation. Plates were centrifuged at 500 × g for 5 min to filter the solution. The plates were fully dried and concentrated using a CentriVap Benchtop Vacuum Concentrator (Labconco, Kansas City, MO, USA). Plates were covered and stored at neg 80°C until ready for LC-MS/MS data acquisition. Samples were resuspended in 200 µL of 50% (v/v) methanol in water. Plates were sonicated for 10 min, followed by centrifugation at 500 × g for 10 min to remove any remaining particulates prior to transfer to a shallow well plate.

### LC-MS/MS data acquisition

LC-MS/MS analysis was performed on a Thermo Scientific™ Q Exactive™ Mass Spectrometer system coupled with a Thermo Scientific™ Vanquish™ UHPLC System (Thermo Scientific, Waltham, MA). Chromatographic separation was performed on a Phenomenex Luna Omega Polar C18 HPLC column (2.1 x 50 mm, 1.6 μm; Torrence, CA), maintained at 40 °C. The mobile phase consisted of H2O (solvent A), and ACN (solvent B), both acidified with 0.1% formic acid. Samples (3 μL) were injected into the column and an initial mobile phase was 5% B and kept for 1.0 min. Mobile phase B was increased in consecutive linear gradients from 5% to 99% over 7.0 min and held for 1.0 min, followed by a linear gradient to 5% B over 0.5 min and held for 0.5 min. It was then increased in consecutive linear gradients back to 99% in 0.5 min and held for 0.5 min, followed by a linear gradient to 5% B over 0.5 min and held for 1.0 min, for a total of 12.5 min. The flow rate was kept stable throughout the run at 500 μL/min. MS analyses for MS1 full scan were carried out using heated electrospray ionization with positive spray voltages of 3.5 kV, sheath gas 53 Arb, auxiliary gas 14 Arb, sweep gas 3 Arb, ion transfer tube temperature 269 °C, aux gas heater temperature 438 °C and Orbitrap resolution of 35,000. The scan range (m/z) was from 150 to 1500. MS/MS spectra were separately acquired using previously described method^49^ with a QC pool sample.

### LC-MS/MS data processing

The raw MS/MS spectra were converted to mzML files using MSconvert (ProteoWizard)^68^ and deposited in GNPS/MassIVE under: MSV000097935. Feature extraction was performed using MZmine 4.2.0 with the following parameters: for mass detection, the factor of lowest signal was 5 for MS1 and 2.5 for MS2. For chromatogram building, the mass tolerance was set as 0.002 m/z or 10 ppm, the minimum consecutive scans as 4, and the minimum height as 5E4. Local minimum search for chromatographic deconvolution was performed with a minimum search range of 0.05 min, minimum ratio of peak top to edge of 1.8, and maximum peak duration of 1.2 min. The peaks were de-isotoped within 5 ppm m/z and 0.05 minutes retention time tolerances, aligned with 0.0015 m/z or 5 ppm mass tolerance and 0.2 minutes retention time tolerance, then gap-filled with 5 ppm m/z tolerance and 0.2 minutes retention time tolerance. Ion forms were assigned using the metaCorrelate module with a retention time tolerance of 0.02 min, minimum feature height of 3E4 and intensity threshold for correlation at 1E4. Feature height correlation was selected using Pearson similarity measure with a minimum correlation of 70 % across a minimum of 3 samples. The feature list was then exported as a feature quantification table (.csv) and an MGF spectra file. Metabolite annotations were performed with the default GNPS Spectral Libraries and the GNPS-BILE-ACID-MODIFICATIONS, GNPS-MASSQL-BILE-ACID-ISOMER, Carnitines_library_2025_testing_V2_MassQL_only and the MULTIPLEX-SYNTHESIS-FILTERED libraries using the Feature-Based Molecular Networking (FBMN) workflow on GNPS2. The spectra were filtered by removing MS/MS fragments within ±17 Da of the precursor m/z and to only keep the six most intense fragments in a ±50 Da window. The spectra were searched against the GNPS libraries with precursor and fragment ion mass tolerances of 0.02 Da and 0.05 Da, respectively, with a cosine score threshold of 0.6 and a minimum of 5 matched peaks. The GNPS2 job is available at: https://gnps2.org/status?task=6ea14530712c4b1ba27f638961b76c24.

### sPLS-R model development

The feature and annotation table were imported into R 4.4.3. Features that elute before 0.7 minutes were removed to avoid dead volume. Total intensity curves were investigated to explore general sample quality. We subtracted features that did not have at least 5 times the peak area compared to blank samples to remove noise. To minimize the confounding effect of inclusion age, we selected a single visit per participant to include in the model, choosing the visit that produced the lowest standard deviation across participants. We applied RCLR conversion using package vegan 2.6-10, after which we applied age as covariate. We removed near-zero variance features. A sparse partial least squares - regression (sPLS-R) model was built using package mixOmics 6.30. Tuning of the model was performed using mixOmics 6.30 and inflection 1.3.7. Variance explained was evaluated using linear modeling, and performance was evaluated using 10-fold cross validation and permutation testing (n = 100). External validation of the model was performed using the visit closest in age to the selected visit for each participant. The same model parameters were applied to generate and evaluate the model for the external dataset. Performance of the model on the external data was assessed using Pearson correlation. Stability of the selected features was evaluated using Jaccard and loading correlation.

### SIRIUS formula prediction

Molecular formula generation and class prediction for the selected features were performed through SIRIUS 6.3.2.^22^ The workflow parameters were set to Orbitrap with 5 ppm MS2 mass accuracy, and [M+H]+, [M+Na]+, [M+K]+, and [M+H3N+H]+ were selected as fallback adducts. The SIRIUS module was run using the default *de novo* + bottom-up strategy for m/z values below 400, with ZODIAC re-ranking. Finally, CSI:FingerID for fingerprint and CANOPUS for compound-class predictions within the score threshold, and CSI:FingerID for structure database search, was set to use PubChem as a fallback with approximate confidence. Compounds with multiple ion charges, a low number of peaks, a mass greater than 850 Da or chimeric spectra were removed from this analysis.

## Statistical analysis

The linear mixed model was performed using package lme4 (version 1.1-37) and lmerTest (version 3.1-3). For selecting features consistently associated with age, we selected for non-sparsity with cutoff <70%. Remaining zero-values were handled using feature-specific pseudocount, defined as half the minimum non-zero intensity for each feature, followed by logL transformation. Age was. Consistency was defined as the absence of detectable inter-individual heterogeneity in age-related slopes.

For detecting which of the features associated with cognitive resilience also changed with age, features were selected for non-sparsity with a cutoff of < 40%. Remaining zero-values were handled using feature-specific pseudocount, defined as half the minimum non-zero intensity for each feature, followed by logL transformation. Centered age was used as predictor, with participant as random intercept and as random slope. Slopes were estimated and p-values were corrected for multiple comparisons using Benjamini-Hochberg false discovery rate.

Statistical analyses for assessing associations between carnitines and cognitive resilience scores were performed in R (version 4.5.2) using a hurdle model (Figure 4a). Step 1 of the hurdle model, assessing whether presence or absence of a metabolite influenced cognitive resilience, was performed using logistic regression. Step 2 was assessing whether, for the samples where a metabolite was detected, abundance of that metabolite was associated with cognitive resilience. This was tested for statistical significance using log-transformed abundance of the metabolite as a linear function of resilience. In both steps, p values were corrected for multiple comparisons using the Benjamini–Hochberg false discovery rate method.

### Mass Spectrometry Search Tool (MASST) searches

We performed batch searches for MS/MS spectra in the GNPS2 repository using MASST tools. Precursor and fragment ion mass tolerance was set at 0.05, minimum cosine threshold at 0.7 and the number of matching fragment ions at 3. For ReDU matching, only scans with at least 2 distinct USI’s were kept for analysis. For the ReDU sample type matches, any type with less than 500 available samples in the Pan-ReDU database was not shown. For microbeMASST, we removed scans that were also detected in human monocultures. Taxonomic assignment was performed with NCBI classifiers, using package taxize 0.10. The sPLS model based on exclusively microbial features was built and evaluated using the same methods and parameters as the original models.

