## Supplemental Information for "Serum metabolic signatures of cognitive resilience in a longitudinal aging cohort"

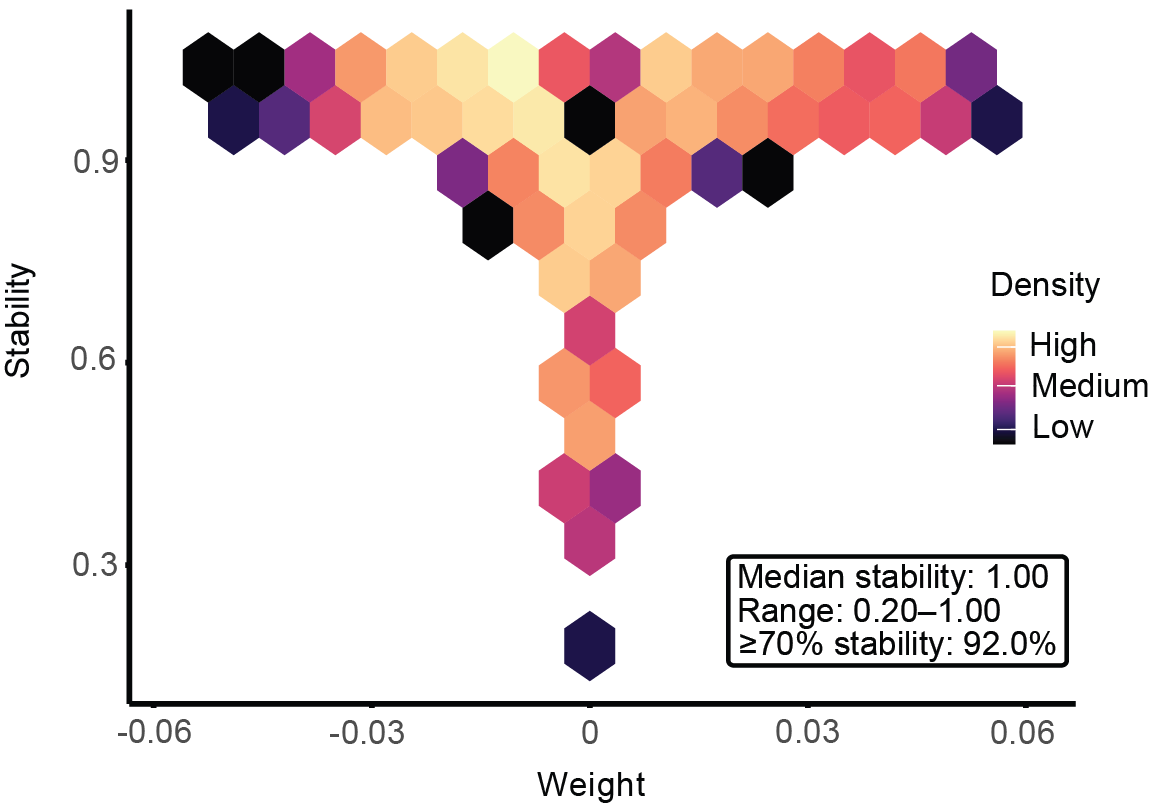

**Supplementary Figure S1.** Stability of selected features across 10-fold cross-validation. A hexplot visualizing the density of features across sPLS weight and selection stability. The inset box summarizes stability metrics.

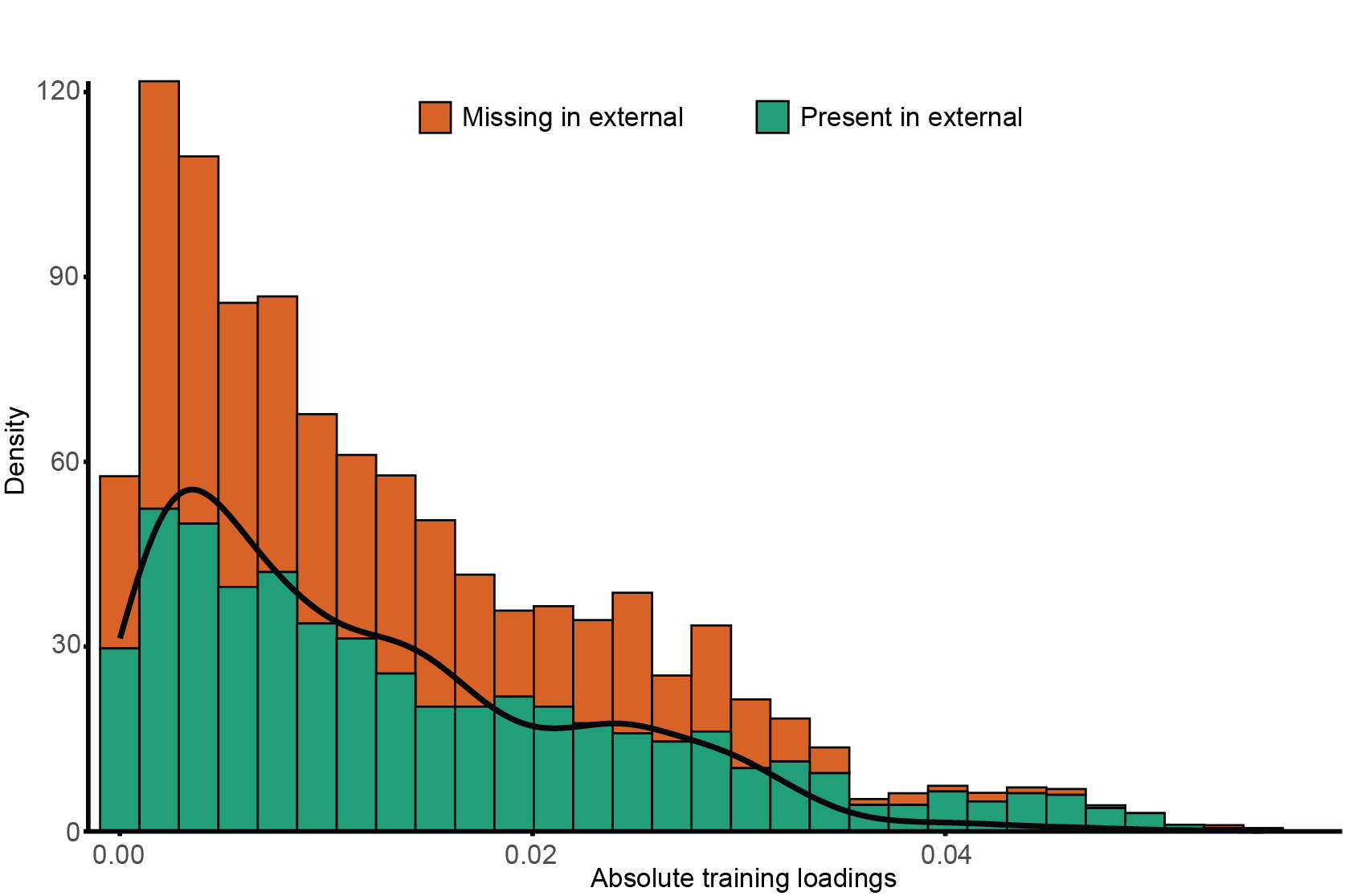

**Supplementary Figure S2.** Externally replicated features across training loadings. A histogram visualizing the distribution of features reselected as important in cognitive resilience in the external dataset. The line visualizes the density of missing features.

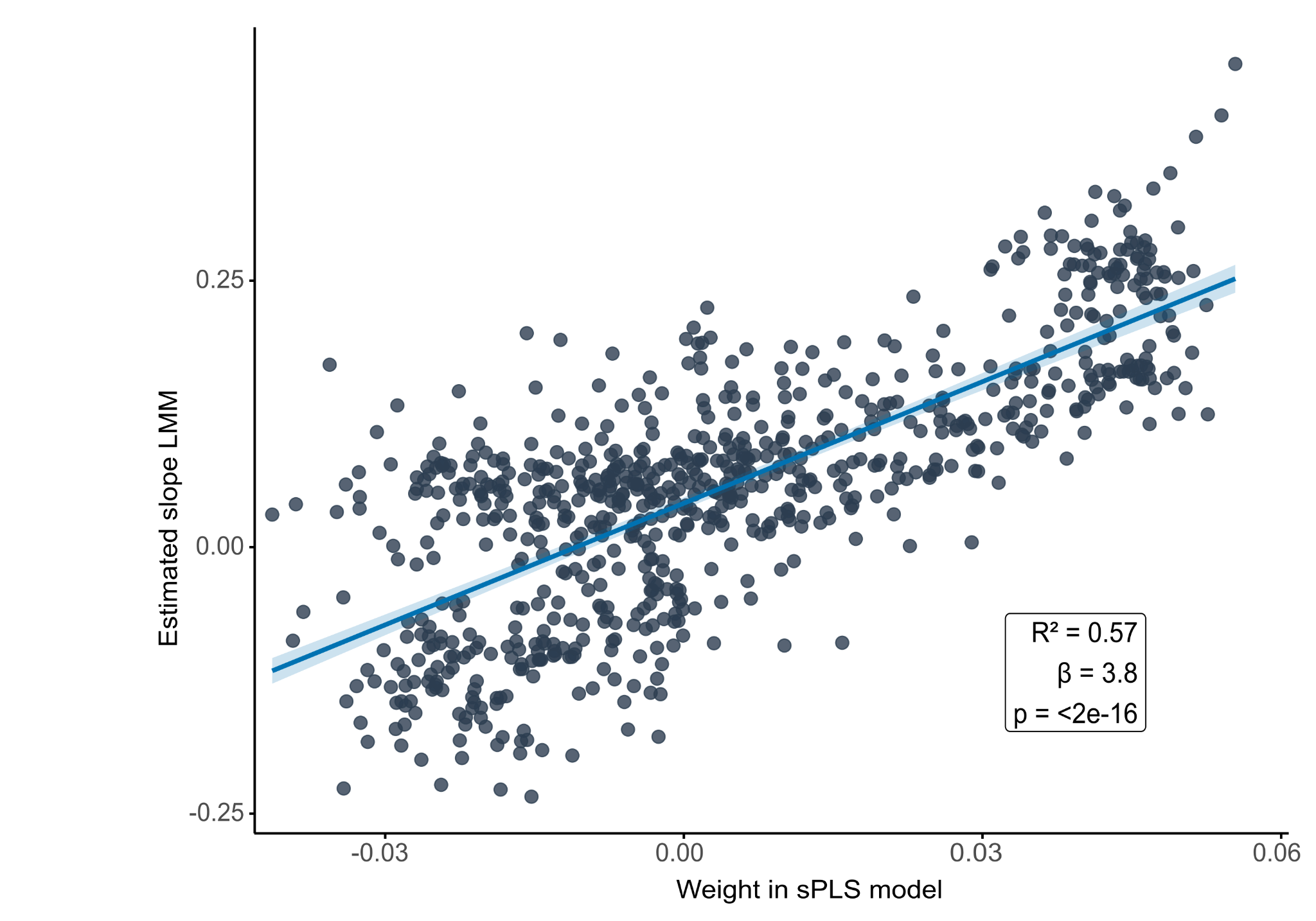

**Supplementary Figure S3.** Association between the weight of features in the sPLS model and their slope across age. Linear regression shows a positive, significant association between weight and slope.

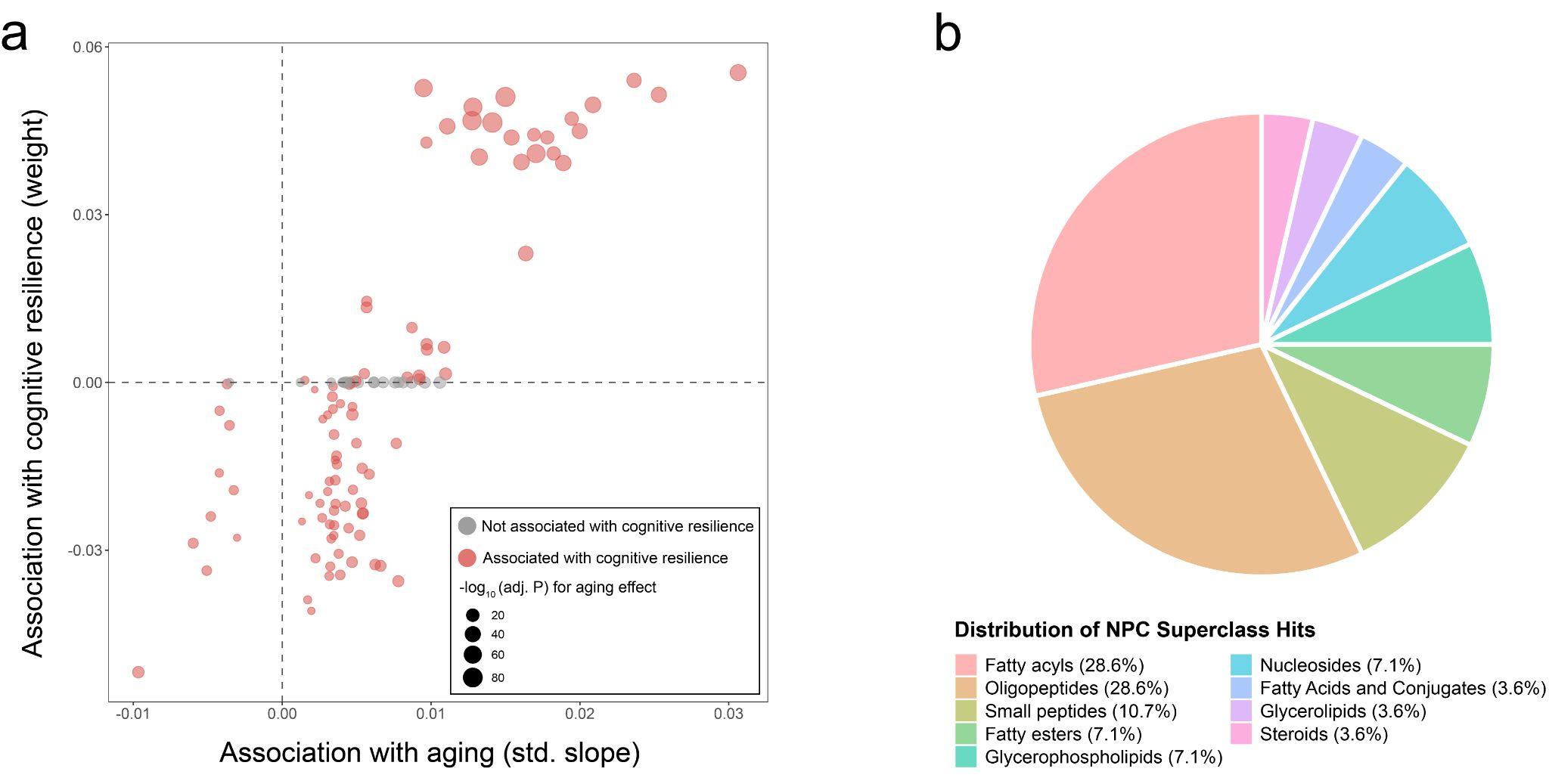

**Supplementary Figure S4.** Association of metabolite features with aging. **a)** Quadrant plot illustrating aging-associated metabolite features and their relationships with cognitive resilience. Red dots represent metabolites associated with aging identified by LMM and with cognitive resilience identified by an sPLS model, whereas gray dots represent metabolites associated with aging only. The size of each dot corresponds to the adjusted p-value for the aging effect. **b)** Distribution of different chemical superclasses within the selected 113 aging-related features, determined by SIRIUS 6.3.2.19.

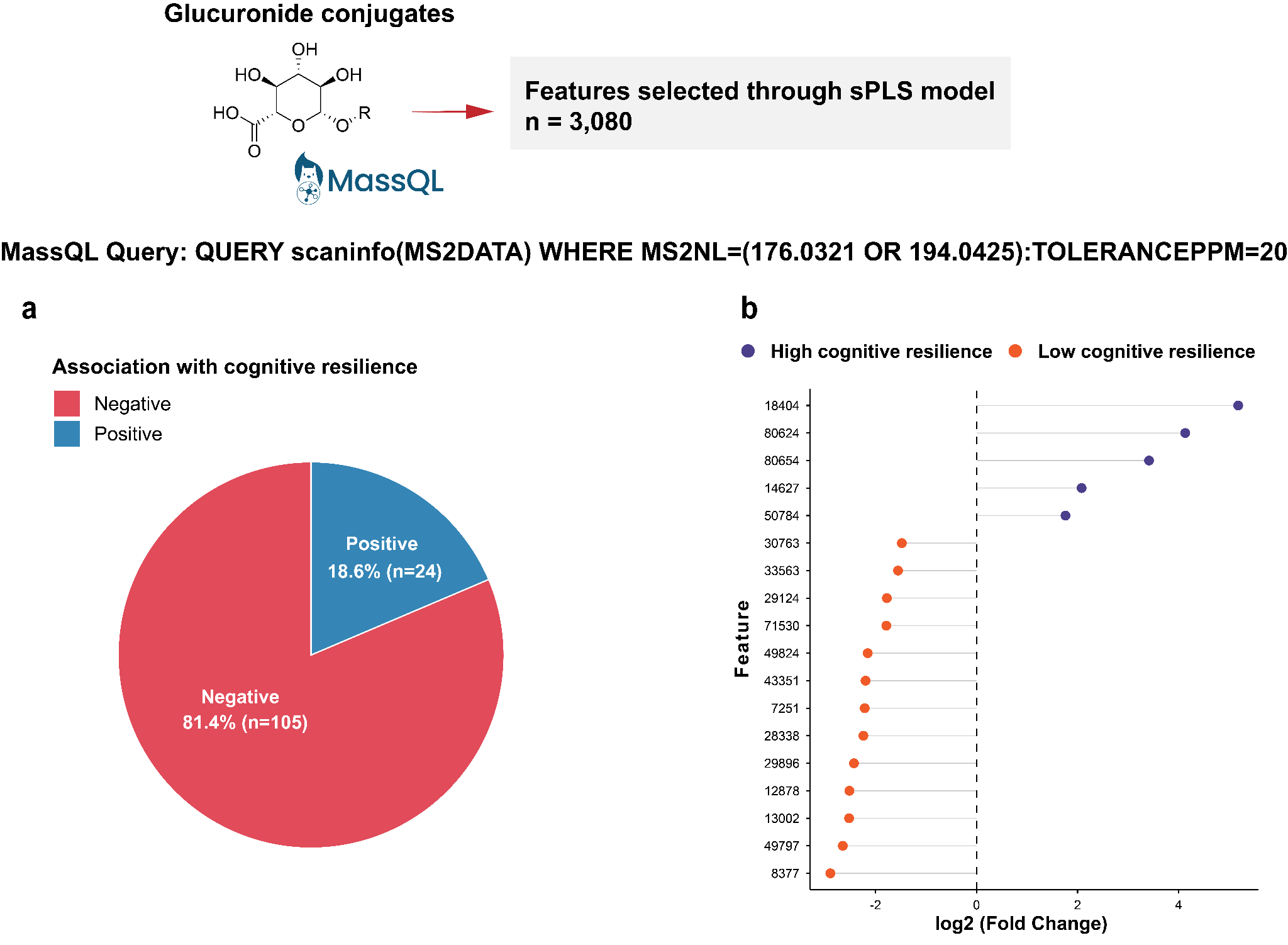

**Supplementary Figure S5.** MassQL analysis revealing associations between potential glucuronide conjugates (among 3,080 features selected through the sPLS model) and cognitive resilience. **a)** Positive or negative associations of retrieved compounds with cognitive resilience based on MassQL results compared with the sPLS analysis. **b)** Fold-change plot showing significantly different peak area distributions (p < 0.05) of retrieved glucuronidated compounds in individuals with high or low cognitive resilience. Statistical differences between high- and low-cognitive-resilience groups were evaluated using the two-sided Mann-Whitney U test, and p values were adjusted for multiple comparisons using the Benjamini-Hochberg method.

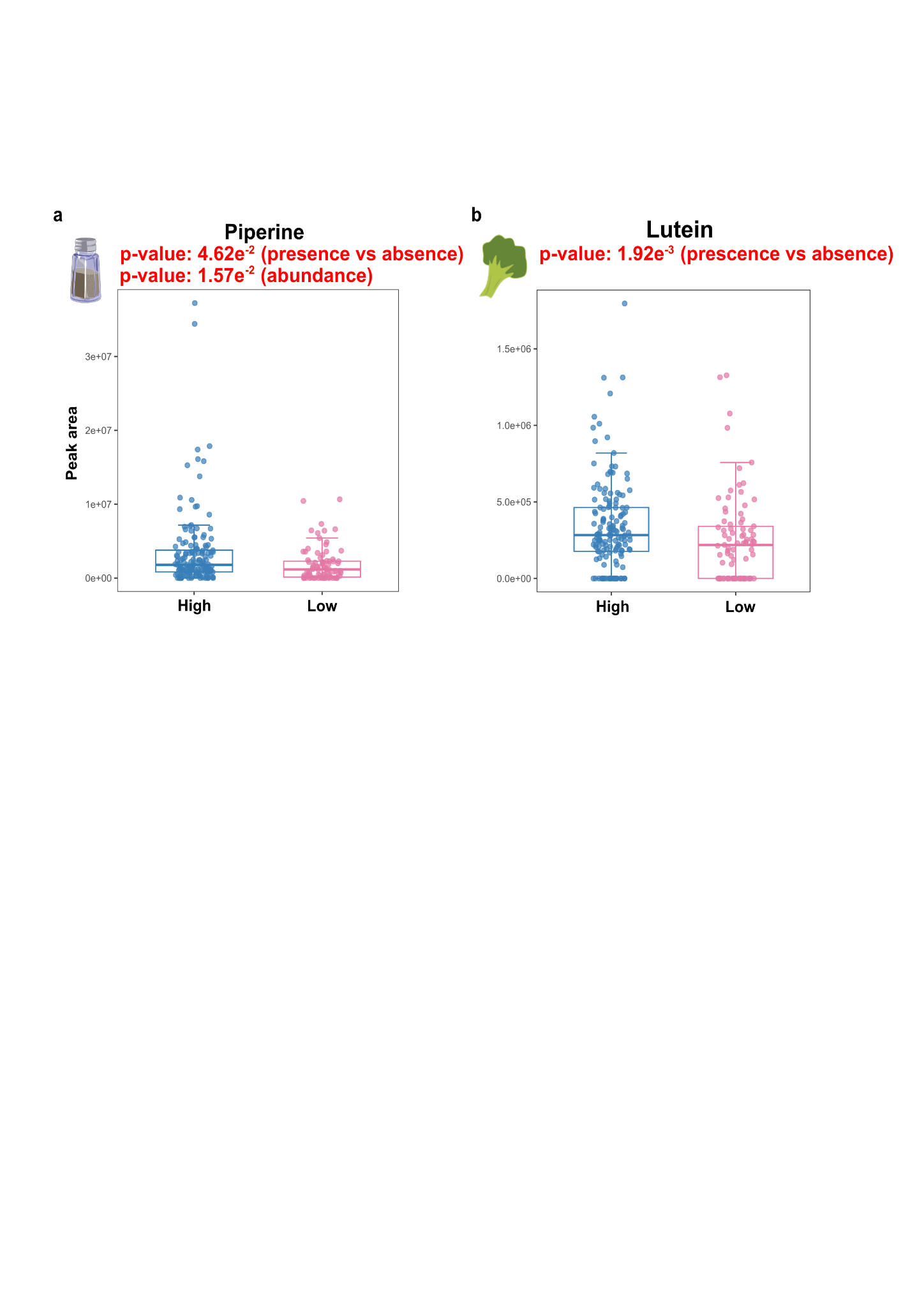

**Supplementary Figure S6.** Peak area distributions of **a)** piperine and **b)** lutein in individuals with high or low cognitive resilience. Statistical differences between high- and low-cognitive-resilience groups were evaluated using the hurdle model, and p-values were adjusted for multiple comparisons using the Benjamini-Hochberg method. Compounds were putatively annotated based on spectral library matching in GNPS2. Icons were obtained from [Bioicons.com](http://bioicons.com).

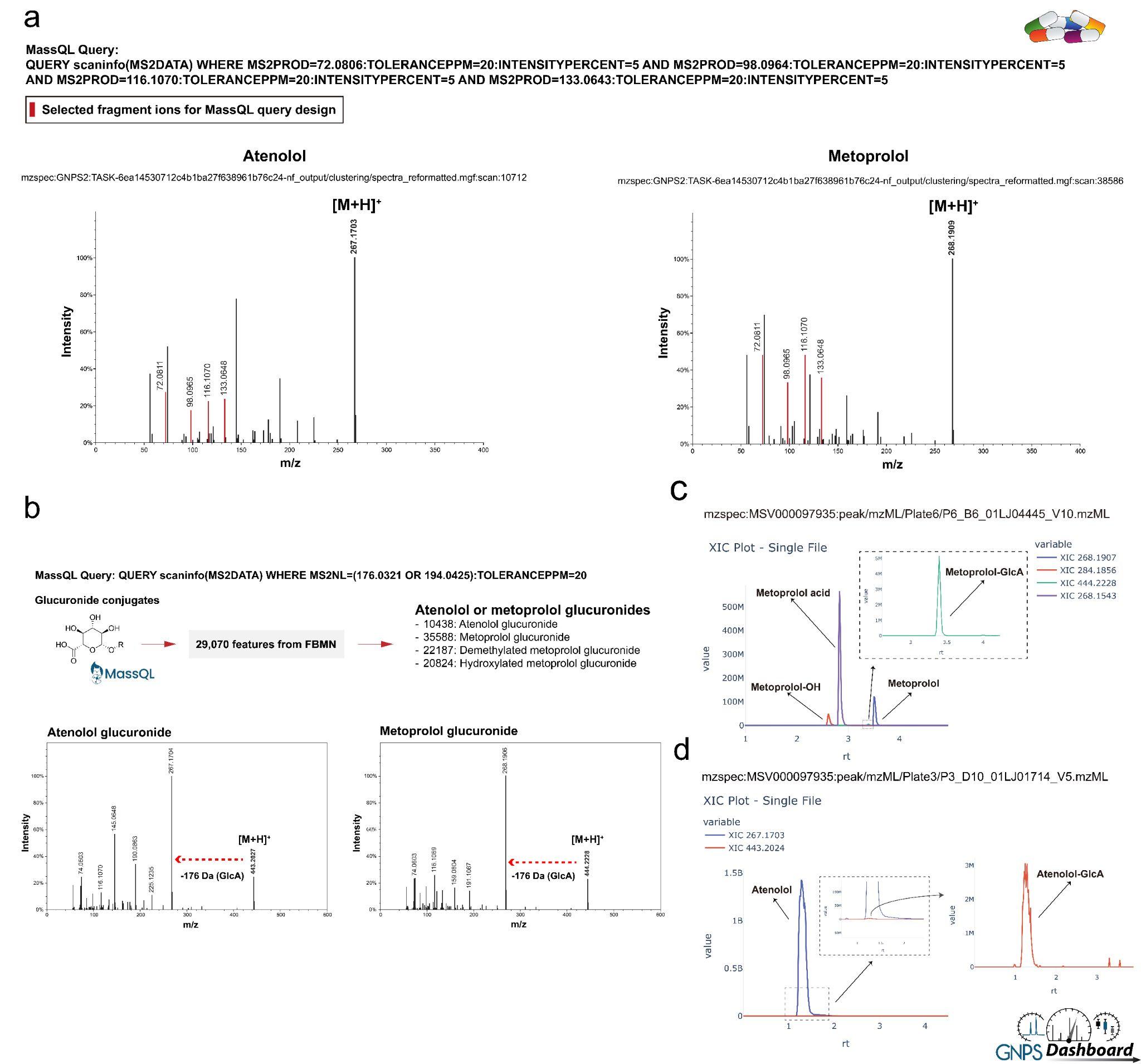

**Supplementary Figure S7.** Designed MassQL queries and mass spectrometric and chromatographic information for atenolol, metoprolol, and their metabolites. **a)** Designed MassQL queries to capture atenolol, metoprolol, and their metabolites, and MS/MS spectra of atenolol (left) and metoprolol (right). **b)** MassQL analysis identifying potential atenolol or metoprolol glucuronides, with MS/MS spectra of the retrieved glucuronides showing the characteristic neutral loss of 176 Da. **c)** Extracted ion chromatograms (EICs) of metoprolol and its metabolites. **d)** EICs of atenolol and its metabolites. MS/MS spectra were obtained through GNPS2, and EICs were generated in the GNPS Dashboard. The icon was obtained from [Bioicons.com](http://bioicons.com)**.**

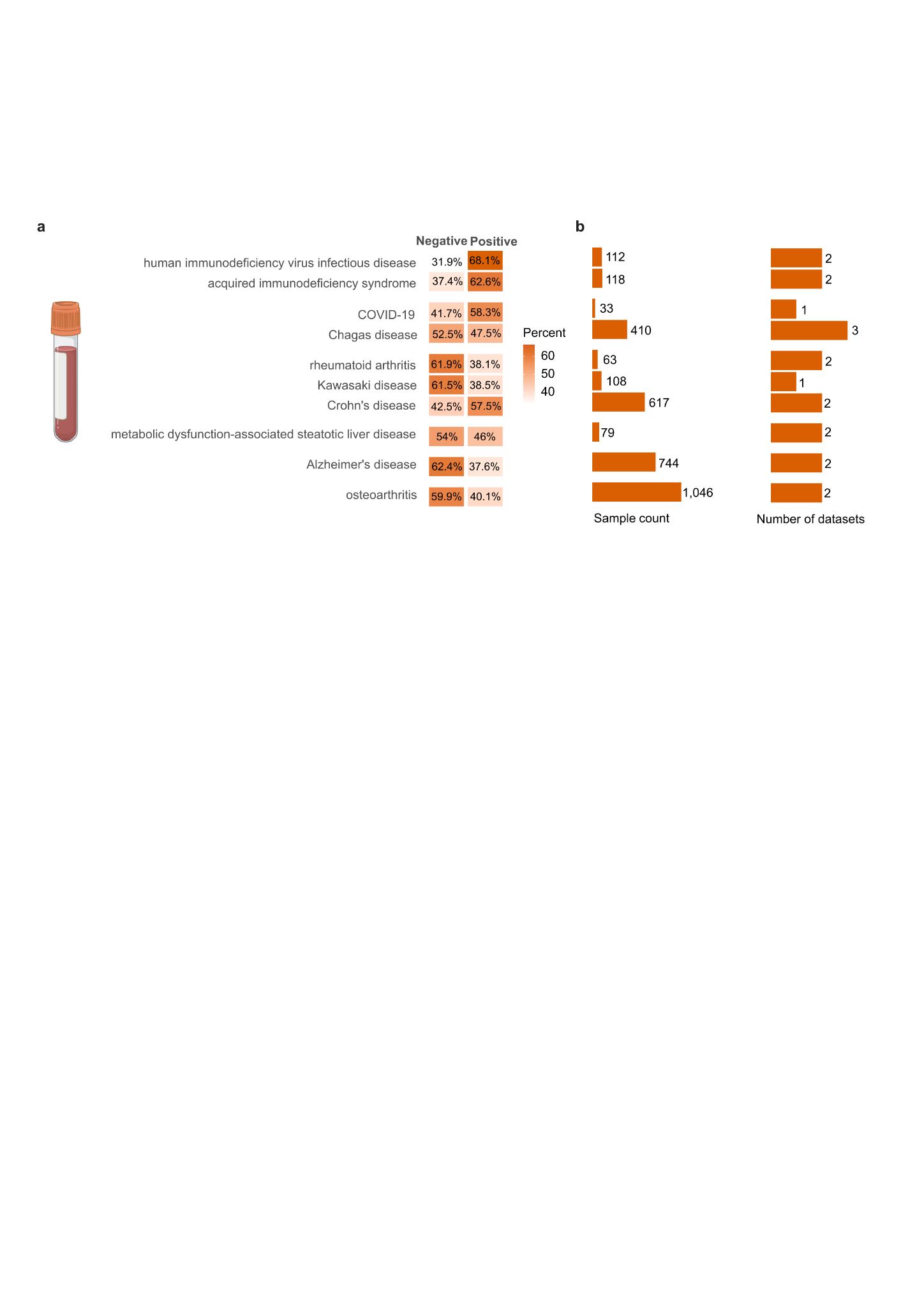

**Supplementary Figure S8.** The repository searches for metabolite features related to cognitive resilience. **a)** Distribution of features matched in human studies in blood samples across different disease states, and **b)** the availability of the total number of datasets in the repository for that disease type.
